# An integrated workflow for structural virology with a 100 keV electron microscope

**DOI:** 10.64898/2025.12.08.693081

**Authors:** Rasangi Pathirage, Moumita Dutta, Ruth J. Parsons, Muralikrishna Lella, Emma Atwood, Qianyi E. Zhang, Aaron May, Alexis Johnson, Xiao Huang, Jordan Flemming, Ujjwal Kumar, Bahjat F. Marayati, M. Ariel Spurrier, Coco Liu, Junlin Zhuo, Ki Song, Radha Devkota Adhikari, Salam Sammour, Victor Ilevbare, Carter Abram, Michael Diaz, Albert Guzman, Jay Rai, Ashwin N. Skelly, Michael P. Hogarty, Kara Anasti, Max Purro, Myles Lindsay, Munir Alam, Drew Weissman, Alon Herschchorn, Beatrice H. Hahn, George M. Shaw, Amit Sharma, Nicholas S. Heaton, Robert J Edwards, Rory Henderson, Thomas Denny, Kevin O. Saunders, Janet Siliciano, Robert Siliciano, Barton F. Haynes, Katarzyna Janowska, Priyamvada Acharya

## Abstract

Cryo-EM has revolutionized structural biology, especially for flexible and heterogeneous samples, although access to high end microscopes that enable these studies remains a bottleneck. While 300 keV microscopes have been the go-to for high-resolution structural determination, they are expensive and restricted to institutional and national facilities needing specialized expertise, with access falling far short of the demand. Here, we present the user-managed operation of a cheaper 100 keV electron microscope within a structural biology laboratory enabling close integration with protein production, biochemical and biophysical studies. We provide details and considerations for the installation of the microscope, its day-to-day maintenance, and operations. Using virus surface glycoproteins as case studies, we illustrate the workflow from grid screening, data collection, and data processing, and provide examples of data quality. This user-administered setup provides a training platform for researchers at all levels, with beginners in cryo-EM achieving proficiency to independently operate the microscope within a month of regular use and training. We have demonstrated routine high-quality low-resolution reconstructions using a Ceta CMOS camera and high-resolution reconstructions enabling building of atomic models using a Falcon C direct detector. While there are several examples of facilities that manage cryo-EM and individual laboratories leveraging cryo-EM, we provide here the first demonstration of a modern group independently doing both successfully, something that has been talked about frequently but rarely seen.

## Introduction

Advances in cryo-electron microscopy (cryo-EM) in the past decade, driven by innovations in sample preparation, microscope optics, detectors, and software for high-throughput data collection and data processing, have firmly established it as the leading technique in structural biology. The development and adoption of direct electron detectors marked a turning point, ushering in the “resolution revolution” (Kuijper et al. 2015). Despite widespread use and accelerated adoption, several bottlenecks remain that hinder the accessibility of cryo-EM for structural determination and have prevented it from fully realizing its true potential. Optimal sample preparation often is a bottleneck and lack of ready access to a screening microscope can slow the pace of sample optimization. Access to high-end microscopes at National Cryo-EM Facilities (Ishemgulova et al. 2023; Haynes 2024; Chiu et al. 2019), has eased access to both screening and data collection needs and has enabled the adoption of cryo-EM by new users through their regular training programs. Nevertheless, geographic distances involving shipping and travelling, as well as the often lengthy wait times for instrument access present substantial hurdles.

While 300 keV microscopes have been the most sought after for high resolution data collection, the potential of cheaper 200 keV and 100 keV microscopes are being realized (Chan et al. 2024; McMullan et al. 2023). Here, we demonstrate the adoption of a 100 keV cryo-TEM (Transmission Electron Microscope) for single particle cryo-EM analysis within a structural biology laboratory, closely integrated with protein production, biochemistry and biophysical studies and enabling an environment where each and every researcher, irrespective of level of expertise in cryo-EM, can expect to become fully proficient in the entire cryo-EM pipeline, starting from sample preparation to data analysis, within a month of regular mentored training. In a recent study, systematic comparisons were performed on proteins of different sizes, to establish a precedent for using a 100 keV TEM for high-resolution protein structure determination (Karia et al. 2025). In this study, our goal is to provide illustrative examples of the data quality achievable with a 100 keV microscope and the impact of the 100 keV pipeline on a structural virology laboratory that drives independent research directions focused on basic science questions relevant for understanding virus entry and evolution, and also provides structural biology support to larger programs for developing preventative and therapeutic countermeausures against viral diseases. We have used virus surface glycoproteins, HIV-1 envelope (Env), Coronavirus spike protein (S) and Henipavirus fusion protein (F) ectodomains to demonstrate data quality in single particle cryo-EM experiments. Further, while the 100 keV Tundra cannot be used to collect tilt series data for tomography, we have used the microscope for visualization of virus surface glycoproteins in situ on viruses and virus-like particles.

## Results

### Site setup and installation

The vision for the 100 keV microscope was for it to be closely integrated within a biochemistry laboratory (**Figure 1A**). Thus, space was identified at the back end of a working protein production and purification laboratory to house the 100 keV TFS Tundra (**Figure 1B**). After identification of the site, the vendor conducted a 24-hour survey that tested for noise, vibration, and electromagnetic fields. After the site passed the survey, construction work began to install electrical and plumbing required for operating the microscope. While the construction work was ongoing, the microscope was shipped. Three crates containing the transformer, chiller, switch, and UPS were delivered to the site earlier, as they needed to be installed ahead of time by the construction team before the installation of the microscope.

**Figure 1:**
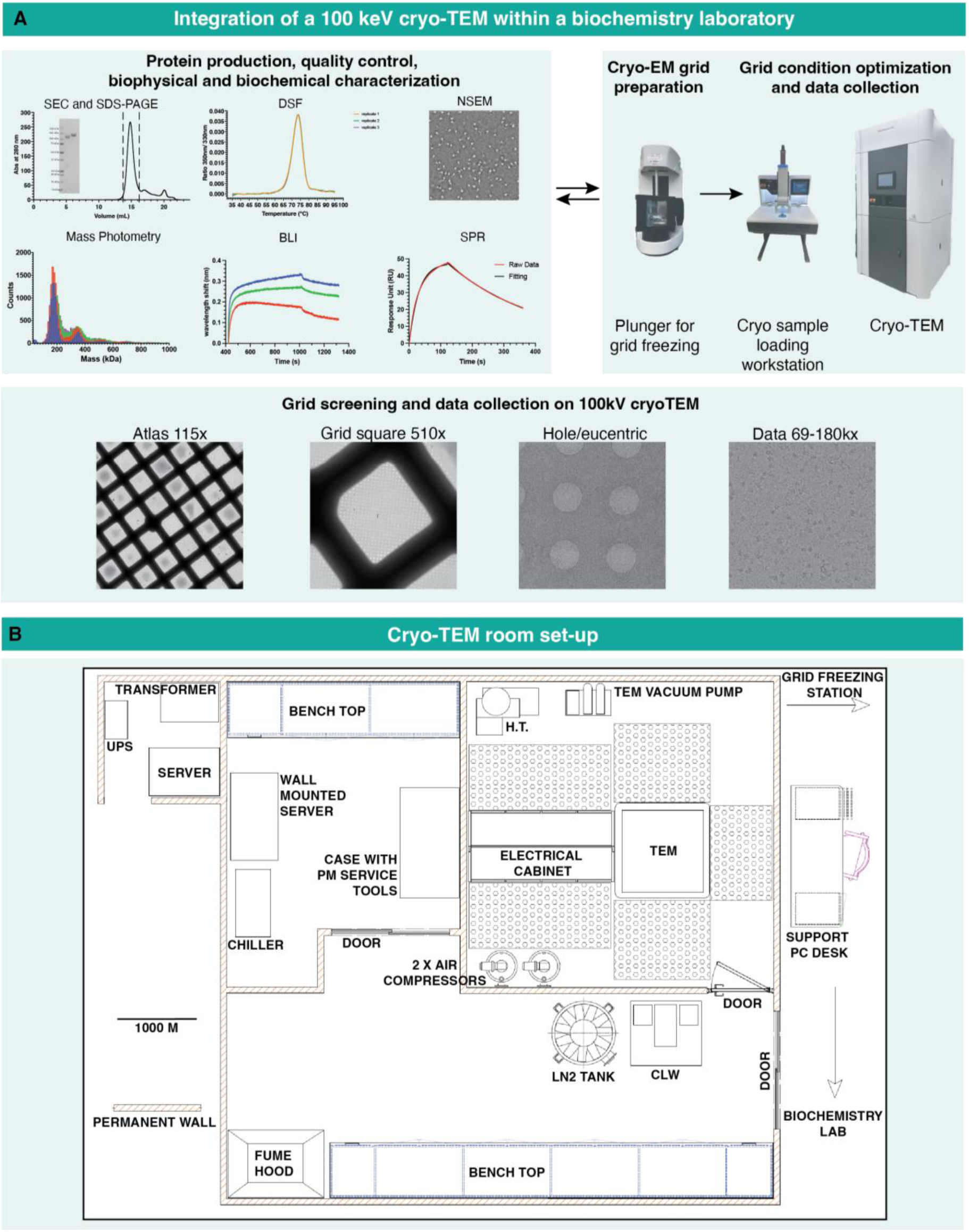
A) Top. Schematic showing the integration of a 100 keV TEM driven cryo-EM infrastructure with protein production, quality control, biophysical and biochemical assays. Bottom. Representative examples of cryo-EM grids viewed at different magnifications. B) Layout of cryo-TEM room.

Installation began with positioning and leveling the console and routing and connecting associated cables and connections, and water hoses. Once the routing of cables, filling the water line, and connecting air to the system were done, an initial powering up of the system was performed and the Field Emission Gun (FEG) was prepared for bake-out. The accelerator and conditioned high tension (HT) were filled with sulfur hexafluoride (SF_6_). Connecting, powering up and deploying the computer server that would control the microscope required coordination between Thermo Fisher Scientific (TFS) and Duke IT teams. Next the loading station was installed and powered up. Initial function checks were performed on the transfer device for optimal loader pick-up position calibration. Sample transfer was tested, warm and cold, between TEM and Loader. After performing alignments and testing the auto-alignment software, the microscope was ready for operations. The microscope was delivered to the Duke warehouse in January 2023. Installation work started May 8^th^, 2023. Installation, testing and sign off were completed on September 6^th^, 2023.

### Training, operations and user management

On-site training was conducted by the vendor on September 7^th^ and 8^th^, 2023. The training was customized to best meet the needs of the user group. The initial training session was attended by 6 team members who had prior data collection experience on the Duke’s Titan Krios microscope. These users became the first group to operate the Tundra, collect data and prepare a Standard Operating Protocol (SOP) that covers microscope operations for screening and data collection. These “superusers” also became the trainers for the “next generation” of Tundra users in the lab. New users perform the screening and data collection setup supervised by the superusers. Once proficient, they are ready to use the Tundra independently and train other users. The current timeline for transitioning from a new user to superuser is ∼ 1 month with regular, consistent practice during this month.

On average 1-5 samples are loaded per day, with the microscope being used both for grid screening and data collection. A typical workday involves a morning session of grid screening, followed by an afternoon session for setting up overnight data collection on prior screened grids. Weekends are reserved for pre-screened samples on which larger datasets need to be collected. The reservations of the microscope for screening and data collection are placed via a shared calendar. Usage is logged on a shared Excel spreadsheet and users communicate via a dedicated Microsoft Teams channel.

### Maintenance of the Microscope

Cryocycle is performed once a week on the cryo-TEM and daily on the Cryo-TEM Loading Workstation (CLW). Users are requested to maintain liquid nitrogen levels of the microscope above 40% all the times, except on days when cryocycle is scheduled. On the day of the cryoocycle, liquid nitrogen levels are maintained at 50-30 %. With 14 fully trained users in the group currently, there is a robust system for ongoing maintenance and training. Microscope maintenance duties are rotated between users ensuring a high level of proficiency among users not only in data collection and data processing but also hands-on microscope operations and management. This management system ensures redundancy, so instead of a dedicated microscopist handling operations and maintenance of the microscope, these responsibilities are shared between users.

The users are trained to troubleshoot any problems encountered during screening, data collection or maintenance of Tundra. However, if needed, further technical assistance is provided by TFS personnel upon request. A support computer is specifically allocated with internet connection and TFS RAPID (Remote Access Program for Interactive Diagnostics) Connection Wizard software installed for remote access (“RAPID Remote Services Connects, security, and use” 2023). The RAPID server is used to build the connection between the vendor (TFS) and user locations without interruptions caused by other networks however the user retains the control of the scope of access by the vendor. Multiple software are used for the daily operations of the Tundra. The hardware-oriented application, Microscope Companion (MiCo) (*Microscope Companion™ 1.16 User Guide* 2025), enabled automated daily alignments and beam optimization with minimal user intervention, with its hardware recovery functions enabling rapid diagnosis and recovery from hardware failures. Optimizations of optics are performed on carbon grids. The AI-powered EPU Software (Bhandari et al. 2025; Thompson et al. 2019) enables automated data acquisition and optimized grid screening. MiCo, Smart EPU, and TEM User Interface software that control microscope functions, are launched and coordinated via the Microscope Software Launcher and Operating System Dashboard Manager (OSD Manager).

### Screening and Data Collection

Screening is performed to assess specimen quality and suitability for data collection. The micrographs are visually inspected for the appearance of intact particles of the expected size. Particle distribution across ice thickness variations is assessed and data collection planned accordingly. The ideal specimen will have an even distribution of intact particles in different orientations. Sometimes an initial assessment of diversity of particle orientation is discernible by visual inspection of the micrograph. Often, however, data processing upto the 2D classification step is necessary to obtain an assessment of particle orientation distribution.

A suitable template for screening is selected out of the three major templates saved as presets in EPU, including the 69 kX, 180 kX and 230 kX magnifications. The lower magnification template 69 kX is used for screening larger particles such as Viruses, Virus Like particles (VLP), Nano Paricles and Liposomes. The most common template used for smaller particles like the Coronavirus spikes, Henipavirus F proteins and the HIV-1 Env is the 180 kX magnification. The 230 kX template (Pixel Size – 1.15 Å, Nyquist resolution – 2.3 Å) is used when the resolution of a certain data set collected on 180 kX is limited due to the nyquist resolution of 180 kX magnification template (Pixel Size - 1.47 Å, Nyquist resolution - 2.94 Å)(Weis and Hagen 2020). The 180kX magnification template is preferred for data collection over the 230 kX magnification template due to the greater field of view with coverage of increased number of particles compared to the 230 kX magnification.

A general workflow for a data collection setup starts with the regular daily alignment followed by the calculation of the electron dose that the sample is being exposed to at data magnification (**Figure 2**). In order to calibrate the shifts at varying magnifications, “Calibrate Image Shifts” is performed from EPU at Atlas (115 X), Square (940 X), Hole/Eucentric (8 kX) and Data Magnifications (180kX/230kX) focused on a sharp feature. Subsequently, the atlas of the entire grid is imaged to select the squares where the data will be collected.

**Figure 2:**
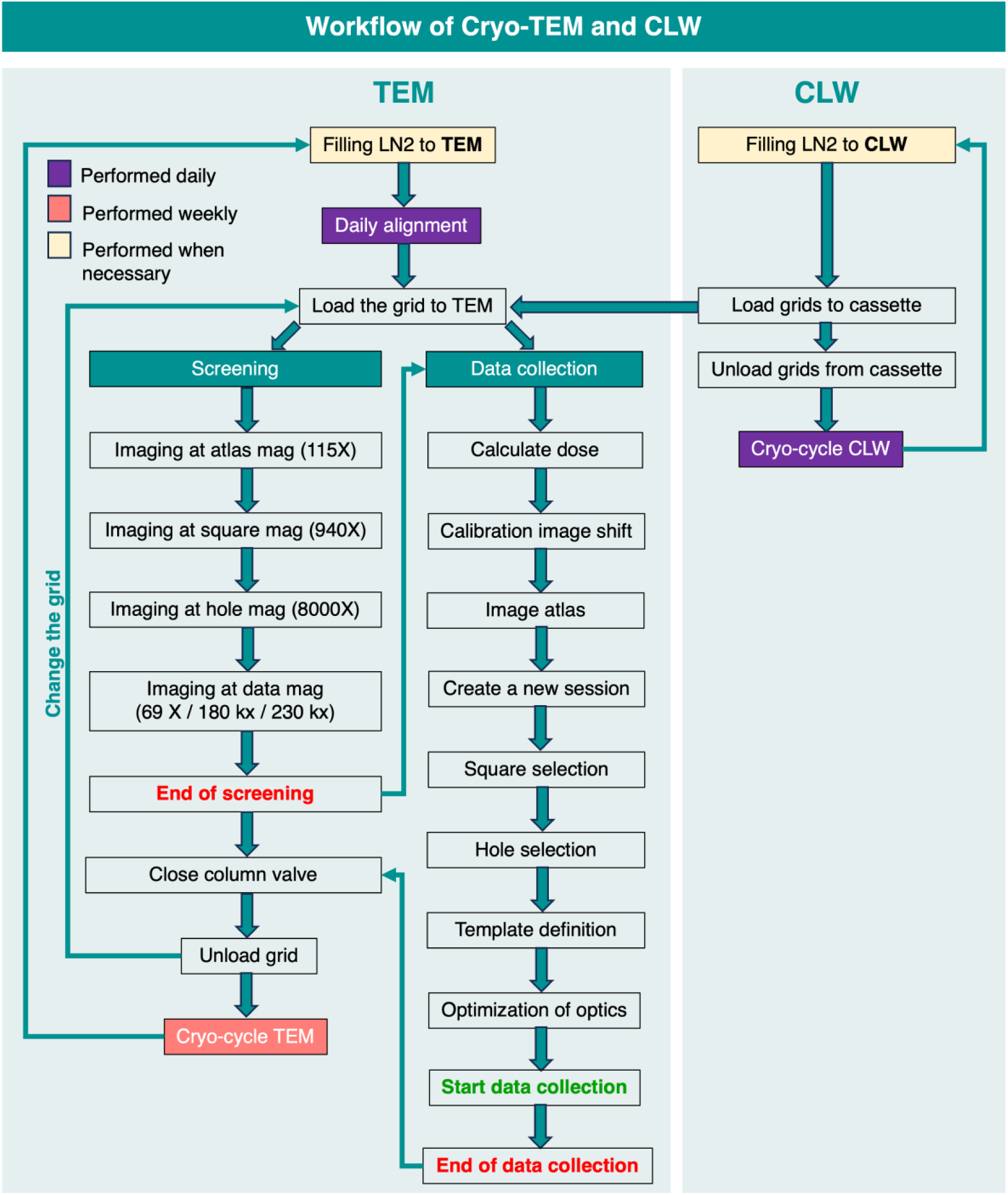
Flowchart showing the workflow of TFS Tundra microscope and CLW for screening and data collection along with the maintenance routine

Our data collection setup leverages the smart selection and filtration options that are available through the EPU software to ensure efficiency of the data collection setup. For large data collections, squares are selected using the smart selection feature, which recognizes and marks squares, color-coding according to the different types of the squares. For short data collections, manual square selection is typically done by the user. The Atlas session is temporarily saved onto the offload data and is deleted post completion. After the square selection, a template is set up for hole selection to identify targets for data collection automatically on each square. The distance between two holes is measured diagonally using EPU to facilitate the auto-identification of target holes within the square. To reduce the need for user intervention for visual identification and removal of unwanted targets, and for increased efficiency of setting up targets, useful options available in EPU such as “Filter Ice Quality” and “Remove Holes Close to Grid Bar” are used. Once satisfied with the template that is used for the target selection of the rest of the squares, a separate template is defined to identify the target area within the hole for data acquisition and the carbon/gold area between the holes for autofocus. The nominal defocus range for the glycoproteins that we work with, including the coronavirus spike, HIV envelope, Henipavirus F protein, is roughly ranging between −0.9 to −3.0 µm. This range is adjusted according to the sample. A short data collection is performed at the beginning to estimate the suitable range of defocus value depending on the contrast between the particles and the background (Cheng et al. 2015).

### Data storage and organization

In parallel with the user managed operations of the Tundra microscope, a user-administered setup for data storage and data processing was established through collaboration with institutional IT experts. Different servers are allocated for the storage of data depending on the stage of data collection and processing. During live data collection, raw micrographs are temporarily stored in a local 80 TB server named “Offload data” (**Figure 3A**). After data collection, files are transferred to a 114 TB “TundraNAS” server. These servers are mounted on 4 workstations that are separately allocated for processing data collected on the Tundra cryo-TEM. Each workstation consists of multiple GPUs (2-4 GPUs) and 40 TB of local storage space. Data processing outputs are saved in the local disk space of each computer. After data processing is complete, the data are transferred to another 58 TB server named “Overflow” for long term storage. After completion of data processing and deposition of the data in the PDB/EMDB, data are transferred to 20 TB external drives for long term storage. Datasets that are no longer needed are permanently deleted from the servers. These include datasets where higher resolution data were obtained from a 300 kV microscope and there is no longer a need to retain the lower resolution data collected on Tundra microscope or instances where binding is not observed in an attempt of getting a structure for a complex of molecules, data will be deleted. All the data processing events are logged in a shared excel spreadsheet which allows real-time determination of available workstations to be used for data processing, monitoring the stage of current data processing at each workstation and easy communication between users.

**Figure 3:**
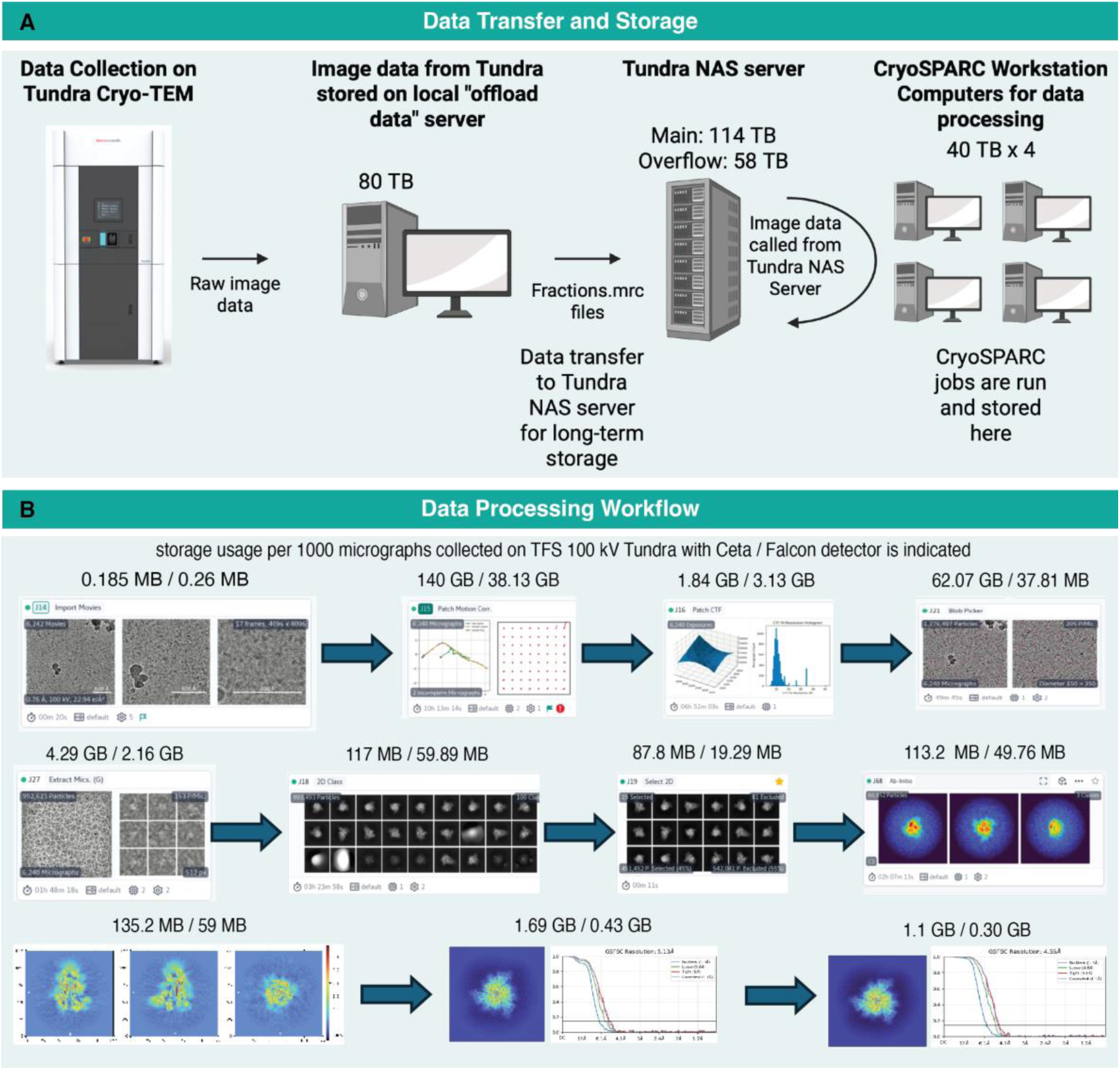
Computational infrastructure for storage and processing of data acquired on the Tundra cryo-TEM A) Schematic showing the storage and data processing setup. B) CryoSPARC workflow with an overnight data collection with Ceta detector is included in the image. The space occupied by each job during processing of data collected with Ceta detector vs Falcon C detector is indicated above each job to get a reasonably high-resolution structure.

While the computational setup for data storage and data processing will likely differ between different laboratories and institutions depending on resource availability, workflow requirements, throughput and other factors, we have described here an active and working setup that serves the needs of multiple projects and users, and has room for adaptive optimizations as needs change and usage increases.

### Data Processing workflow and storage of CryoSPARC jobs

A general workflow for processing the data collected on 100 keV Tundra microscope to generate and refine cryo-EM density maps is illustrated in Figure 3B. While different software suites may be used for data processing, our current setup uses CryoSPARC (Punjani et al. 2017). After “Importing” the movies, Patch Motion correction and Patch CTF estimation are performed to correct beam induced motions, phase flipping and amplitude. Blob picker and extraction from micrographs are performed to pick particles using the inputs of particle size and suitable box size, respectively. Particles are down sampled 4-fold to reduce storage and run time for efficient data processing. Multiple rounds of 2D classification are performed to eliminate junk particles. Ab-initio reconstruction is performed and reconstructions are generated *de novo* from the clean stack of particles. The ab initio generated volumes are used as input for heterogeneous refinement jobs. Instances where prior map information are available, volumes low pass filtered to a resolution of 20 Å may be used as the initial model for the heterogeneous refinements. After multiple rounds of heterogenous refinements accompanied with removal of additional junk particles, particles are re-extracted to the original box size (Punjani et al. 2017). Non-uniform refinements, local refinements and homogenous refinements are performed to further refine the maps. Adjustments are made to this typical processing workflow as needed. While Blobpicker is our first choice for picking particles, Template-based picking and Topaz (implemented within cryosprac) are also sometimes used.

Data storage is a key consideration, not only for the raw data collected but also the files generated during data processing. Figure 3B lists data storage requirements at each stage of our workflow for data collected on Tundra with Ceta F CMOS detector vs Falcon C direct electron detector (DED). The data that were collected with the Ceta detector are in MRC format whereas after installing the Falcon C detector the micrographs are stored and processed as EER format files. The MRC format files store the information based on pixel intensities of exposure fractions, whereas the EER format stores the information as x, y and time coordinates of electron detection on sensor (Guo et al. 2020). The amount of storage occupied by each CryoSPARC job for processing data collected on Falcon C detector has reduced compared to the data collected on Ceta detector.

### Structural determination of Coronavirus spike proteins using 100 keV cryo-TEM

To assess data quality, we examined cryo-EM maps of spike (S) ectodomains of the human Coronavirus OC43 (HCoV-OC43), one of the viruses responsible for common cold (Wang et al. 2022) **(Figure 4** and **Table 1)**. While the samples were prepared aiming to determine structures in complex with antibodies (see methods for details), only ligand-free spike particles were observed in the cryo-EM datasets. We examined the refined maps of the ligand-free S-OC43 to assess map quality and resolutions that can be achieved on this sample using the 100 keV Tundra cryo-TEM, with either the Ceta or the Falcon camera and how this compares with data collected on the 300 keV Titan Krios.

**Figure 4.**
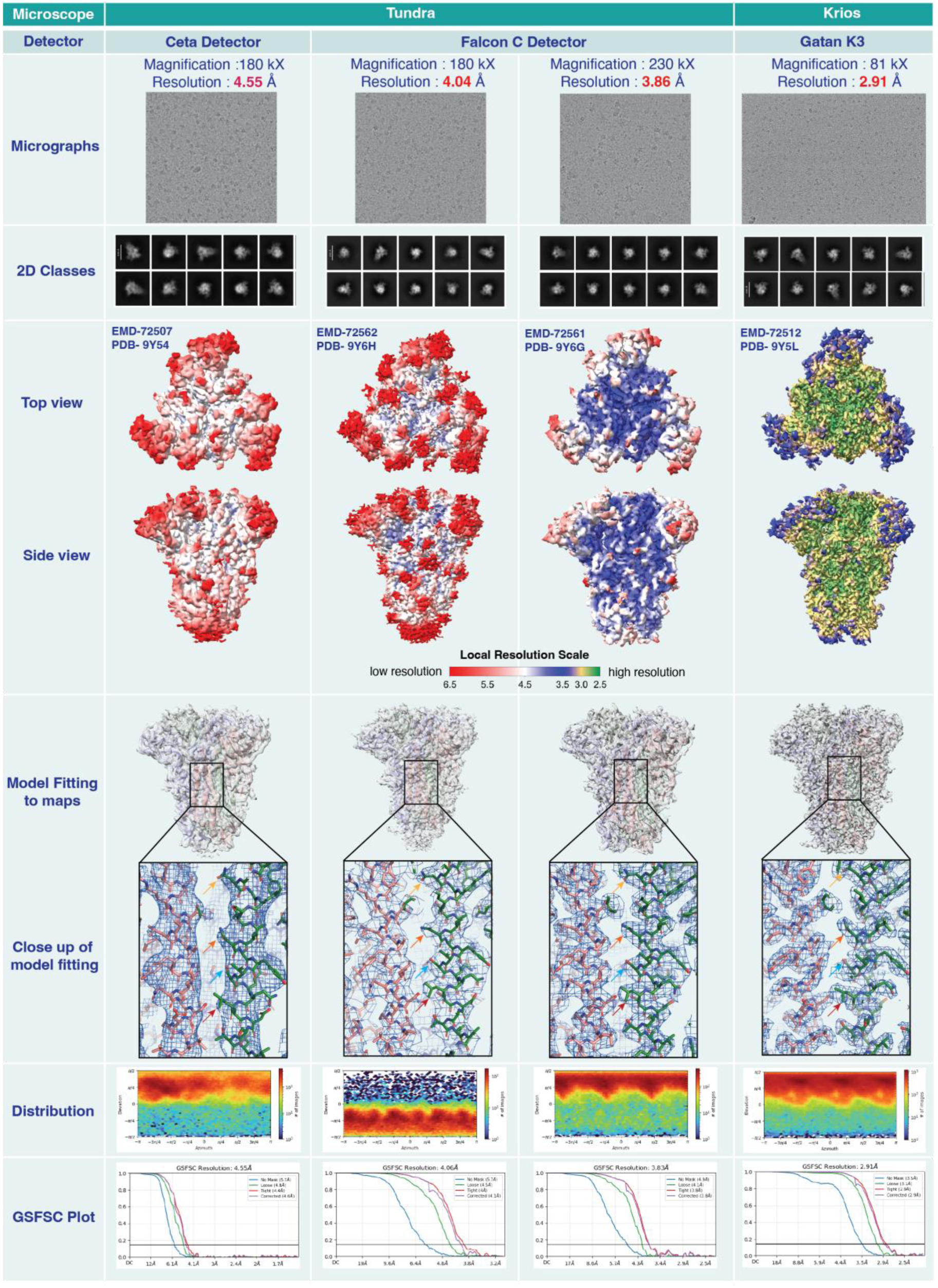
Comparative study of HCoV-OC43 spike structures generated from Ceta and Falcon C detectors in the 100kV Cryo-TEM with a side-by-side comparison with high resolution structure obtained from 300kV Cryo-TEM of the same spike.

**Table 1:** Cryo-EM data collection and refinement statistics for OC43 spike.

| Microscope | Tundra |  |  | Krios |
| --- | --- | --- | --- | --- |
| Detector | Ceta Detector | Falcon C Detector |  | Gatan K3 |
| Accelerating voltage | 100 keV | 100 keV | 100 keV | 300 keV |
| Magnification | 180,000 X | 180,000 X | 230,000 X | 81,000 X |
| Pixel size | 0.76 Å | 1.5 Å | 1.15 Å | 1.1 Å |
| Electron exposure | 26.75 e/Å<sup>2</sup> | 31.10 e/Å<sup>2</sup> | 30.93 e/Å<sup>2</sup> | 64.50 e/Å<sup>2</sup> |
| Exposure time | 1.25 s | 6.75 s | 3.93 s | 2.8 s |
| Defocus range | -3 to -0.9 | -3 to -1.5 | -3 to -0.8 | -2.8 to -1.5 |
| Image size | 4096 x 4096 | 2048 x 2048 | 2048 x 2048 | 4096 x 5760 |
| Number of micrographs used for data processing | 6278 | 7675 | 10,347 | 5985 |
| Box size for extraction from micrographs | 400 px | 256 px | 300 px | 320 px |
| Symmetry imposed | C3 | C3 | C3 | C3 |
| Map resolution | 4.55 Å | 4.06 Å | 3.83 Å | 2.91 Å |
| FSC threshold | 0.143 | 0.143 | 0.143 | 0.143 |
| Number of particle images | 195, 202 | 97,429 | 242,516 | 660,143 |
| Concentration of Spike | 2 mg/mL | 1.7 mg/mL | 1.7 mg/mL | 1.5 mg/mL |

Data collected on the Tundra with the Ceta detector yielded a map with a global resolution of 4.55 Å with imposed C3 symmetry (**Table 1**). After the Falcon C detector was installed, two data sets were collected at 2 different magnifications - 180 kX and 230 kX, yielding maps with of 4.06 Å and 3.86 Å, respectively. Focusing on the central helices for visualizing local resolution features, the map obtained using data collected on Ceta detector showed separation of the helices but the map was tubular with little to no side chain definition. The maps obtained from data collected with the Falcon C camera were better resolved with clearly visible side chain densities. While the resolution could be further improved with data collected on the ligand-free spike on the 300 keV Titan Krios, the quality of the maps obtained using the Falcon C camera was sufficient to allow building of an atomic model.

Similar results were obtained with the spike protein from the spike proteins of two other Coronavirus: HKU1, a coronavirus that causes upper respiratory disease in humans with symptoms of the common cold, but can advance to pneumonia and bronchiolitis (McCallum et al. 2024; Esper et al. 2006; Woo et al. 2005), and a bat coronavirus, PRD0038 that has potential to spillover to humans (Feng et al. 2023). Using the Tundra/Falcon C setup, a 3D reconstruction of global resolution 3.72 Å was obtained for both HKU1 spike (C3 symmetry imposed) and PRD0038 spike (**Figure 5** and **Table 2**).

**Figure 5.**
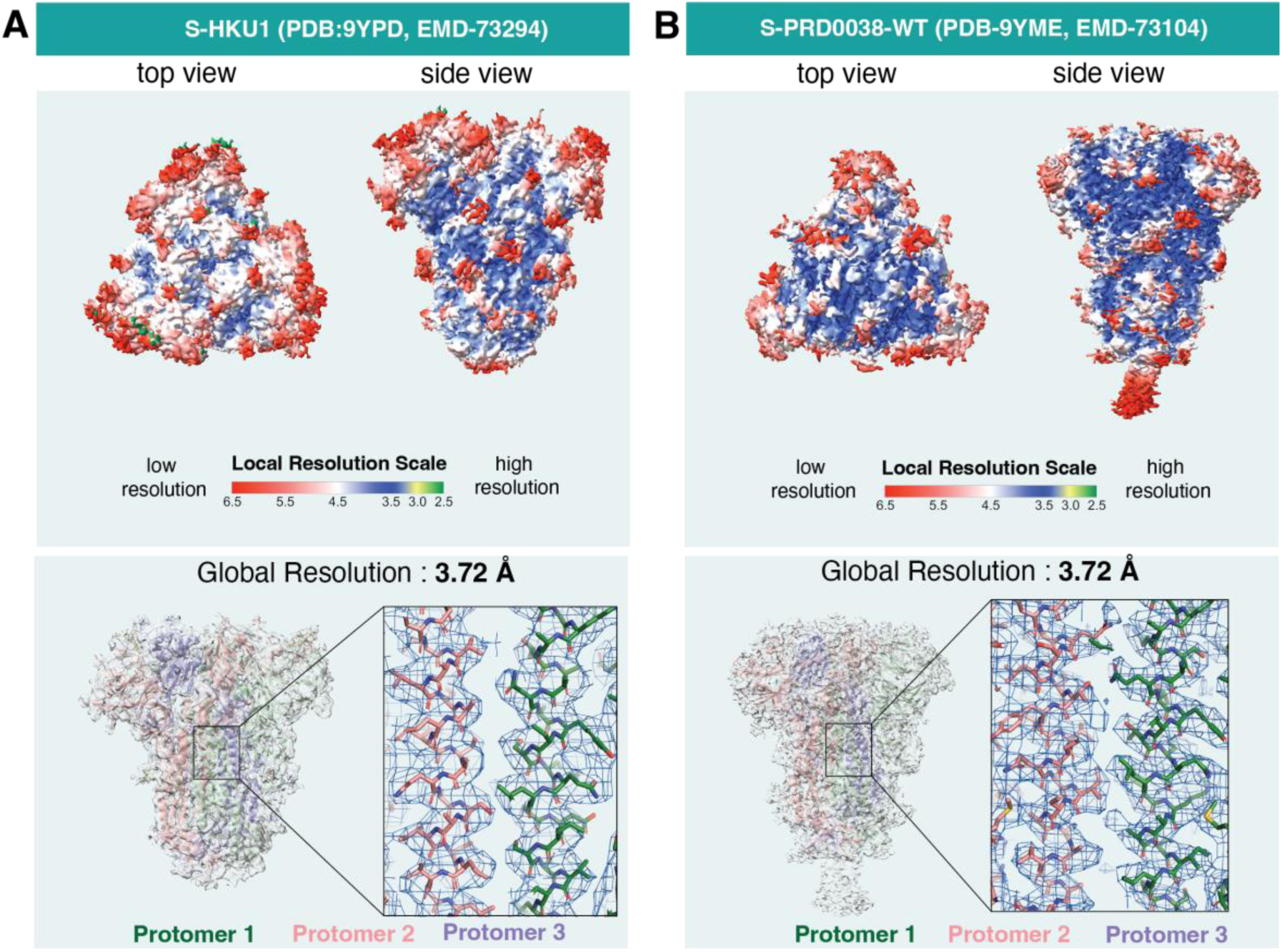
Structures of spike proteins obtained from 100 keV Tundra with Falcon C detector, **A.** The ligand free structure of the endemic coronavirus HKU1 spike structure **B.** The structure of PRD0038 spike. The upper panel shows the local resolution distribution of both maps. The lower panel shows the overall fitting of the models to maps and the closeup of the central helices region of the spikes highlighting the distinctive features of the Cry-EM maps for side chains at respective resolutions.

**Table 2:** Cryo-EM data collection and refinement statistics for spike proteins from HKU1 and PRD0038.

| Microscope | Tundra |  |
| --- | --- | --- |
| Detector | Falcon C Detector |  |
| Sample | S-HKU1 | S-PRD0038 |
| Accelerating voltage | 100 keV | 100 keV |
| Magnification | 180,000 X | 180,000 X |
| Pixel size | 1.47 Å | 1.47 Å |
| Electron exposure | 29.17 e/Å<sup>2</sup> | 31.00 e/Å<sup>2</sup> |
| Exposure time | 6.75 s | 6.75 s |
| Defocus range | -3.0 to -0.9 | -2.5 to -0.5 |
| Image size | 2048 x 2048 | 2048 x 2048 |
| Number of micrographs used for data processing | 11247 | 10659 |
| Box size for extraction from micrographs | 256 px | 256 px |
| Symmetry imposed | C3 | C1 |
| Map resolution | 3.72 Å | 3.72 Å |
| FSC threshold | 0.143 | 0.143 |
| Number of particle images | 149,190 | 715,662 |
| Concentration of Spike | 1.46 mg/mL | 2 mg/mL |

**Table 3:** Cryo-EM data collection and refinement statistics for HIV-1 Env samples.

| Microscope | Tundra |  | Krios |
| --- | --- | --- | --- |
| Detector | Falcon C Detector |  | Gatan K3 |
| Sample | HIV-1 Env with vaccine elicited antibody | Unliganded HIV-1 Env | Unliganded HIV-1 Env |
| Accelerating voltage | 100 keV | 100 keV | 300 keV |
| Magnification | 180,000 X | 180,000 X | 81,000 X |
| Pixel size | 1.47 Å | 1.47 Å | 1.08 Å |
| Electron exposure | 30.00 e/Å<sup>2</sup> | 30.45 e/Å<sup>2</sup> | 61.65 e/Å<sup>2</sup> |
| Exposure time | 6.75 s | 6.75 s | 2.8 s |
| Defocus range | -3.0 to -0.9 | -3.0 to -0.9 | -3.0 to -0.8 |
| Image size | 2048 x 2048 | 2048 x 2048 | 4096 x 5760 |
| Number of micrographs used for data processing | 13541 | 8077 | 17700 |
| Box size for extraction from micrographs | 256 px | 256 px | 320 px |
| Symmetry imposed | C1 | C1 | C1 |
| Map resolution | 4.64 Å | 6.74 Å | 4.00 Å |
| FSC threshold | 0.143 | 0.143 | 0.143 |
| Number of particle images | 364,320 | 134,111 | 888,825 |
| Concentration of sample | 1 mg/mL | 4.44 mg/mL | 4.44 mg/mL |

**Table 4:** Cryo-EM data collection and refinement statistics for Henipavirus F protein samples.

| Microscope | Tundra |  |  |  |
| --- | --- | --- | --- | --- |
| Detector | Ceta Detector |  |  | Falcon C Detector |
| Sample | AngV-F | AngV-F hexamer | HepV-alpa2-F | LayV-F with antibody |
| Accelerating voltage | 100 keV | 100 keV | 100 keV | 100 keV |
| Magnification | 180,000 X | 180,000 X | 180,000 X | 180,000 X |
| Pixel size | 0.76 Å | 0.76 Å | 0.76 Å | 1.47 Å |
| Electron exposure | 24.63 e/Å<sup>2</sup> | 24.63 e/Å<sup>2</sup> | 27.42 e/Å<sup>2</sup> | 30.25 e/Å<sup>2</sup> |
| Exposure time | 1.25 s | 1.25 s | 1.25 s | 6.75 s |
| Defocus range | -2.8 to -1.7 | -2.8 to -1.7 | -3.0 to -1.7 | -2.5 to -0.5 |
| Image size | 4096 x 4096 | 4096 x 4096 | 4096 x 4096 | 2048 x 2048 |
| Number of micrographs used for data processing | 2189 | 2189 | 2189 | 3508 |
| Box size for extraction from micrographs | 512 px | 720 px | 512 px | 256 px |
| Symmetry imposed | C1 | C1 | C1 | C3 |
| Map resolution | 7.5 Å | 17.13 Å | 7.5 Å | 4.46 Å |
| FSC threshold | 0.143 | 0.143 | 0.143 | 0.143 |
| Number of particle images | 108,592 | 7,092 | 108,592 | 245,019 |
| Concentration of sample | 1.5 mg/mL | 1.5 mg/mL | 1.5 mg/mL | 1.5 mg/mL LayV-F with Fab in 1: 4.87 ratio |

Taken together, these results demonstrate the ability of a 100 keV microscope to deliver high quality maps for Coronavirus spike proteins that allow atomic level interpretation.

### Structural determination of HIV-1 envelope (Env) using 100 keV cryo-TEM

Compared to CoV spike ectodomains, single particle cryo-EM of HIV-1 Env samples is typically more difficult, owing to the higher conformational heterogeneity, fragility, and tendency of the sample to adhere to air-water interfaces resulting in preferred orientation (Parthasarathy et al. 2024). Thus, while there are numerous cryo-EM structures of ligand-free CoV spikes (Tortorici et al. 2019; Li et al. 2019; Jin et al. 2025; Walls et al. 2016; Zhang et al. 2024; Stalls et al. 2022), structures of ligand-free HIV-1 Env are relatively rare (Liu et al. 2008; Parthasarathy et al. 2024; Wrapp et al. 2023).

We next illustrate the data quality of HIV-1 Env samples that can be achieved on a 100 keV cryo-TEM (**Figure 6**). With the Ceta camera the resolutions of maps obtained from HIV-1 Env samples, ligand-free or bound to antibodies elicited naturally or by vaccination, were limited to 8-12 Å, which allowed visualization of subunit interfaces and bound antibodies (**Figure 6A**), but without any resolution of secondary structures or side chains.

**Figure 6.**
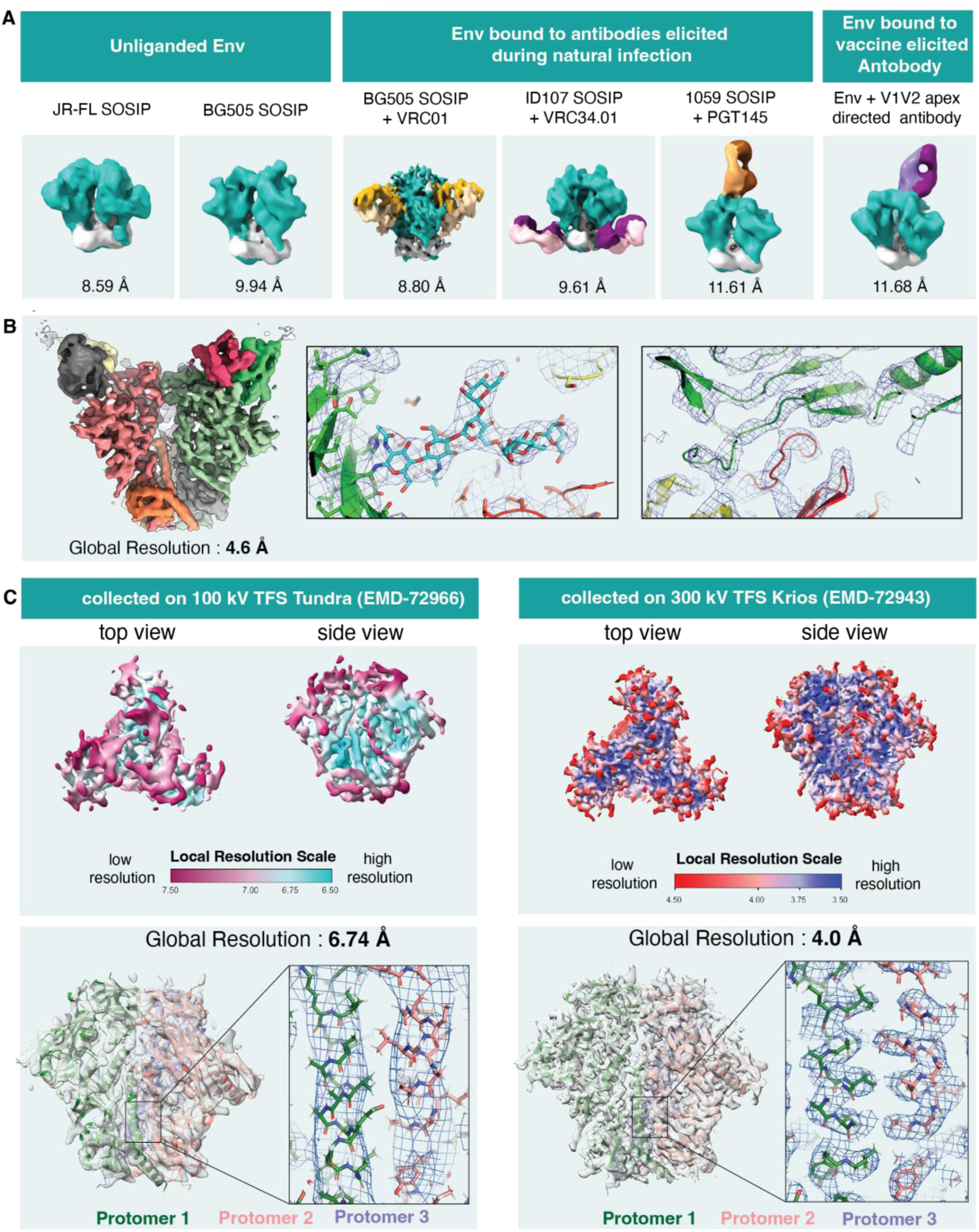
Structures of HIV-1 Env obtained from 100 keV Tundra. **A.** Gallery of HIV-1 Env cryo-EM density maps obtained using Ceta camera. Structures of ligand-free Env from JR-FL and BG505 isolates and of Env bound to antibodies elicited during natural infection or by vaccination are shown. **B.** Cryo-EM map of HIV-1 Env bound to a vaccine elicited antibody obtained using a Falcon C camera. Left. Cryo-EM map of the complex colored by chain. Right. Zoomed-in views with the map shown as a blue mesh and the underlying fitted model shown in cartoon representation. A glycan at the Env/antibody interface is shown in stick representation (middle panel). **C.** Comparison of ligand-free HIV Env BG505 SOSIP.Q653L structures obtained from TFS 100 keV Tundra with Falcon C detector (left) versus 300 keV Krios (right). The local resolution distribution of each map is displayed followed by a panel showing the model fitting. The closeup of the central region of the HIV envelope illustrates the difference between the maps with distinctive features for side chain fitting in the map generated from the Krios data.

With data collected on the Falcon C camera, we were able to substantially improve map resolution with a structure of an Env bound to a vaccine elicited V3-glycan supersite targeting antibody yielding a resolution of 4.6 Å, with clearly resolved antibody-Env interface, including well-defined density for interfacing glycans (**Figure 6B**).

A lignd-free Env sample yielded a map with a global resolution of 6.7 Å, with well-resolved gp41 alpha helices but without any side chain definition (**Figure 6C**). For the same sample, a map with a global resolution of 4.0 Å and well-resolved side chains that allowed unambiguous atomic level model building could be obtained with data collected on a 300 keV Titan Krios.

Thus, although the map resolutions of the HIV-1 Env samples were overall poorer than what we were able to obtain for CoV spike samples, substantial improvements were noted for data obtained using a Falcon C camera versus a Ceta C-MOS camera. The 100 keV Tundra/Falcon C setup can be used for determining structures of antibody-bound Env with maps resolved well enough to allow atomic level model building. For ligand-free Envs, this setup would allow assessment of sample quality and suitability for obtaining a high quality 3D reconstruction, but data from a 300 keV microscope would be needed for atomic level analysis and for more advanced analysis, such as assessment of conformational variability and heterogeneity from the cryo-EM data (Parthasarathy et al. 2024).

### Structural determination of Paramyxovirus surface glycoproteins using 100 keV cryo-TEM

The Paramyxovirus family includes Henipaviruses such as Nipah that cause yearly outbreaks of deadly disease, as well as highly infectious viruses such as measles and mumps (May and Acharya 2024). Two glycoproteins on the surface of Paramyxoviruses, an attachment protein that binds host receptor and a fusion protein that mediates virus and host membrane fusion, together facilitate entry of virus into host. Due to their essential functions in the Paramyxovirus life cycle and their surface location, these glycoproteins are targets for the development of preventive and therapeutic countermeasures.

Using the 100 keV Tundra/Ceta camera setup, we determined the structure of the Angavokeley (AngV) fusion (F) protein, revealing novel features such as a stable, well-defined stalk and an enlarged central cavity, features that were later verified and resolved further with a cryo-EM dataset collected on a 300 keV Titan Krios (May et al. 2025) (**Figure 7A**). From the same dataset, we also obtained structures of AngV-F protein multimers. In particular, a hexameric arrangement of the F proteins was observed that demonstrates the ability of the F protein ectodomains to self-associate and may have implications for how the native F proteins associate on the virus surface. Evidence of F ectodomain association was also observed for other strains, such as the Hendra virus.

**Figure 7:**
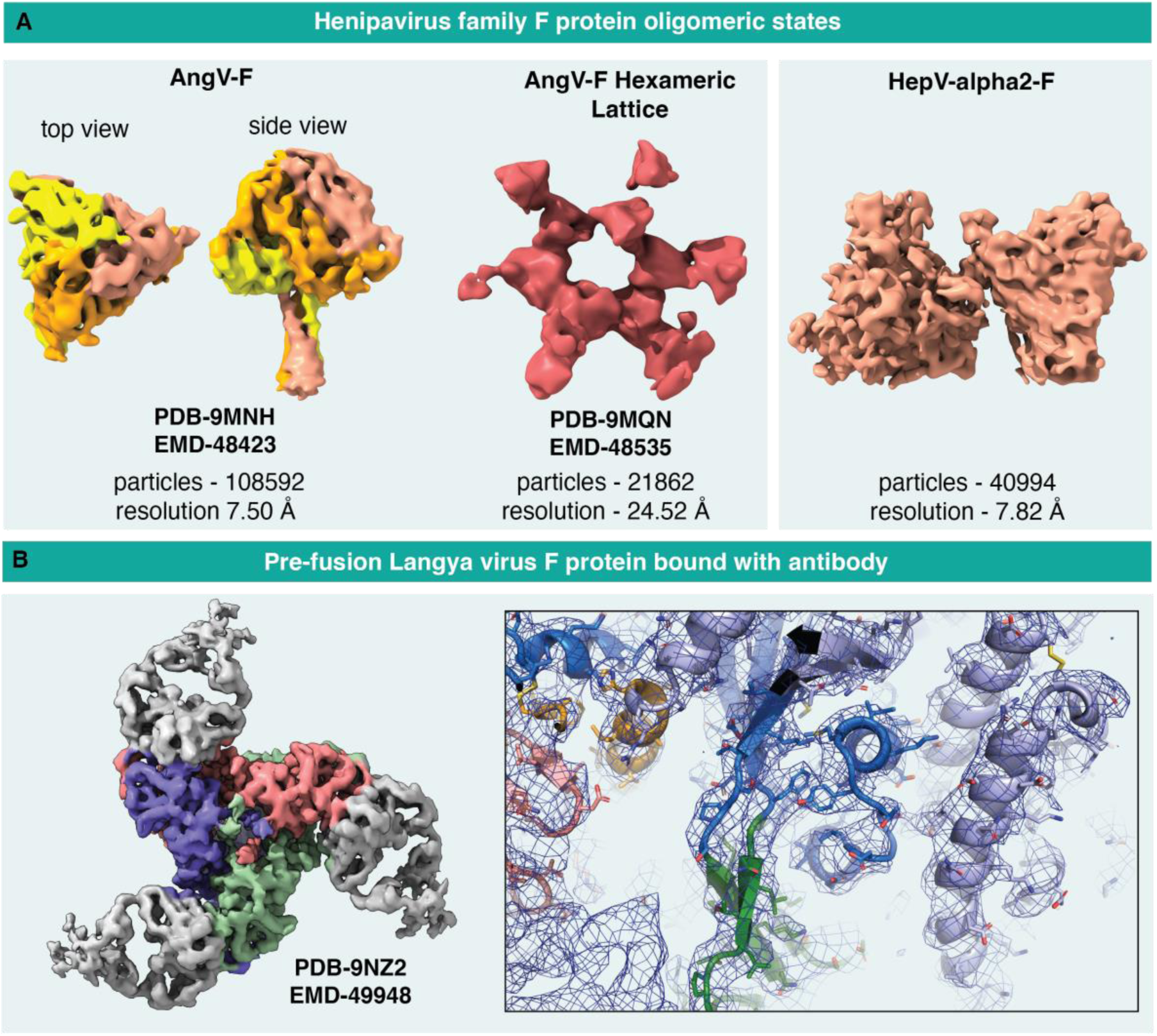
Structures of Henipavirus glycoproteins obtained from 100 keV Tundra **A.** Refined cryo-EM maps of Henipavirus fusion proteins obtained from data collected using the Ceta camera. From left to right; AngV F ectodomain trimer; hexameric arrangement of AngV-F ectodomain obtained from the same protein preparation and the same dataset; and dimer-of-trimers of Hendravirus alpha2 -F protein;. **B.** Refined cryo-EM maps of antibody-bound pre-fusion Langya virus F protein obtained from data collected using the Falcon C camera. Left. Cryo-EM map showing the F protein colored by chain and the bound antibody in grey. Zoomed in image with cryo-EM map shown as a blue mesh and underlying fitted model in cartoon and stick representation.

While useful information could be obtained on the ligand-free F proteins using the Tundra/Ceta setup, the resolutions were limited to visualization of domains. These datasets provided opportunity for robust sample optimization prior to the collection of data on the Titan Krios, which led to high resolution structures that allowed atomic level model building (May et al. 2025)

Collecting data with the Falcon C detector on a pre-fusion Langya virus F protein (LayV-F) bound to a vaccine elicited antibody yielded a 4.46 Å global resolution map that allowed atomic level model building with high quality definition of the antibody-F protein interface (**Figure 7B**) (May et al. 2025).

### Visualizing viruses, virus-like particles, nanoparticles and liposomes using 100 keV cryo-TEM

While single-particle cryo-EM provides detailed structural analysis of purified proteins and complexes by computationally averaging thousands of identical particles, studying the architecture of supramolecular assemblies such as viruses, virus-like particles (VLPs), nanoparticles and liposomes requires 3D reconstruction using cryo-electron tomography (cryo-ET), which involves imaging the specimen at various tilt angles and reconstructing a 3D volume by combining the images from the tilt angles. Although, the Tundra microscope is not equipped to collect cryo-ET data as the sample stage cannot be tilted, we have used it effectively to visualize liposomes, virus and virus-like particles **(Figure 8)**.

**Figure 8:**
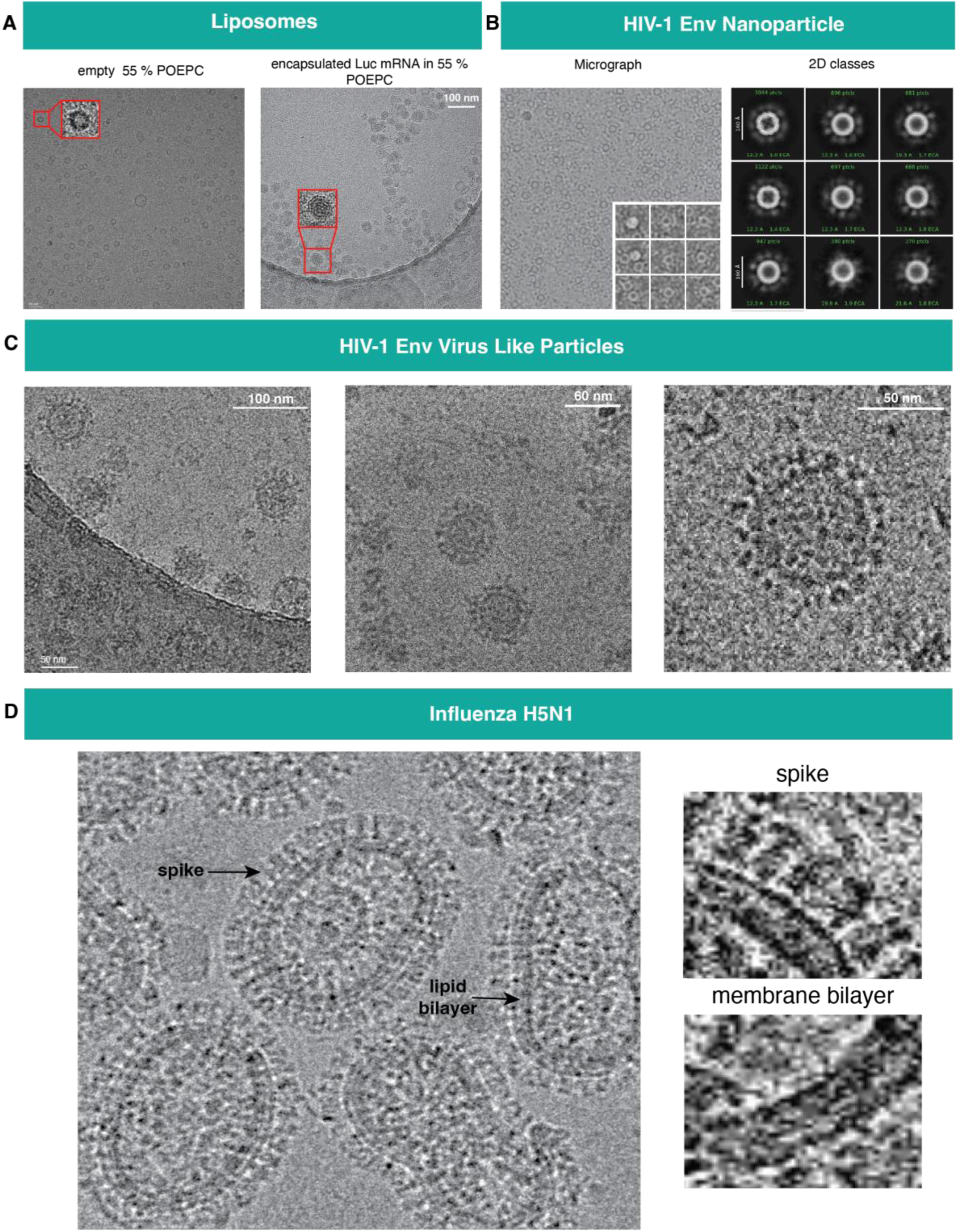
Visualization of supramolecular protein assemblies. A) Left. empty liposomes formulated with 55 % POEPC. Right. liposomes formulated with 55 % POEPC encapsulated with Luc mRNA, imaged with Falcon C detector B) Nanoparticles decorated with an HIV-1 Env-based immunogen on the surface. The first image shows a micrograph of the particle images with Falcon C detector and the inset shows particles extracted from micrographs, the second image shows representative 2D classes. C) Virus Like Particles (VLP) studded with HIV Env, micrographs obtained at different magnifications highlights the details of the particles. First image of the panel was taken at 69 kX magnification with Ceta detector, second at 180 kX magnification with Falcon C detector and third is a zoomed-in view of a VLP imaged at 69 kX magnification with Falcon C detector. D) Left. H5N1 influenza viruses imaged with Ceta detector, Right. zoomed-in views of glycoprotein spikes (top) and membrane bilayer (bottom).

Lipid nanoparticles and liposomes are routinely used as carriers for delivering cargo (Xu et al. 2022; Thirumal, Aravindhan, and Ac 2023). Here we assess the integrity and distribution of liposomes in a 55% 1-palmitoyl-2-oleoylphosphatidylcholine (POEPC) formulation, by vitrifying the liposome preparation and imaging it using the Tundra/Ceta setup **(Figure 8A)**. Well dispersed liposome particles were observed. We also imaged a sample of the liposome preparation with mRNA. The empty liposome that appeared as a dark circle enclosing a lighter inner empty space was clearly distinguishable from the mRNA encapsulated liposome where the entire liposome was uniformly dark.

Engineering multimerization of vaccine immunogens enables better crosslinking of B cell receptors (Ingale et al. 2016). Two commonly used platforms for multimeric presentation of immunogens are self-assembling nanoparticles and virus-like particles (VLPs) that are engineered to display multiple copies of immunogens on their outer surface. Figure 8B and 8C show examples of a ferritin nanoparticle and a VLP preparation, respectively with HIV-1 Env displayed on the surface.

Imaging an influenza specimen revealed expected pleomorphism of the virus particles, with the membrane bilayer clearly discernible and a dense array of glycoproteins visible on the surface **(Figure 8D)**.

Thus, imaging on the 100 keV Tundra microscope is a rapid method for assessing size distribution and dispersity of a liposome (or lipid nanoparticle preparation) and for ascertaining whether it is empty or filled with cargo. The ability to clearly visualize distinct surface glycoproteins on viruses and VLPs provides a robust method for screening and optimizing sample preparation for subsequent cryo-ET data collection.

## Discussion

Surface glycoproteins play an essential role in the life cycle of viruses. They bind host receptors and facilitate fusion of virus and host membranes to mediate virus entry into host cells. Due to their exposed location on the surface of viruses, these glycoproteins are targets for host antibodies, and the focus of preventative and therapeutic countermeasures development. Structural biology of viral surface glycoproteins has played a central role in the development of vaccines and therapies (Derking and Sanders 2021; Byrne and McLellan 2022). Prior to the widespread adoption of cryo-EM, due to their glycosylated and flexible nature, determining structures of these proteins by X-ray crystallography often required protein engineering and biochemical modifications, and the bottleneck of obtaining well ordered crystals for structure determination slowed progress (Fernandez-Leiro and Scheres 2016). Improvements in cryo-EM structural determination spurred by the “resolution revolution” changed this and resulted in increasingly facile and rapid structure determination without the need for extensive modifications such as loop truncations and deglycosylation (Fernandez-Leiro and Scheres 2016).

The realization of the power of atomic level structural determination in accelerating both basic and translational research has led to an ever increasing demand for resources. A particularly challenging bottleneck has been the availability of high end microscopes suitable for generating data that allow atomic level model building. While 300 keV microscopes are preferred for single particle cryo-EM analysis, there has been increasing adoption of the generally less expensive 200 keV and 100 keV microscopes (McMullan et al. 2023; Karia et al. 2025). Yet, these have so far required facility level maintenance requiring a dedicated staff for operations and management. In this study, we have demonstrated the integration of a 100 keV cryo-TEM within a structural virology laboratory, and through case studies have illustrated its use for the study of virus surface glycoproteins. Among the factors that have enabled the successful user-managed operations described here are a motivated and talented workforce, reliance on standardized protocols and robust SOPs, and close collaboration and responsiveness of the vendor that ensures minimum downtime. Additionally, the implementation of AI-enabled automated alignment protocols has greatly facilitated the onboarding and training of new users.

The transition from a Ceta to a Falcon C camera was instrumental for obtaining 3D reconstructions suitable for atomic level model building. For all instances tested, data collected on a 300 keV microscope yielded better resolution compared to the 100 keV microscope. The resolutions that could be achieved from the 100 keV Tundra cryo-TEM was dependent on sample type. Coronavirus spike samples consistently yielded the highest quality reconstructions suitable for building high quality atomic models with side chain definition. For HIV-1 Env and Henipavirus F protein samples, although useful information could be obtained from the ligand-free glycoproteins, their antibody-bound forms yielded reconstructions suitable for atomic model building with side-chain level resolution at the antigen-antibody interface including resolution of interfacial glycans. Thus, while the data collected on a 300 keV microscope is likely to be superior to the 100 keV data, in many instances, maps generated from data collected on a 100 keV microscope are sufficient to provide the information needed. In general, antibody-bound complexes fared better than ligand-free glycoproteins.

In summary, we have demonstrated here how a 100 keV cryo-TEM can be leveraged for accelerating structural virology, impacting the speed and throughput of discovery both in basic research on the structure and conformations of virus surface glycoproteins, as well as in resolving antibody epitopes at high resolutions to provide valuable information for vaccine design pipelines. Continued optimization of the microscope hardware and the software to enable more robust operations may reduce maintenance costs in the future, enabling widespread adoption.

## Acknowledgements

This work was supported by the following NIH grants: U54 AI170752, R01 AI145687, R01 AI165947 and R01 AI 165147 to P.A., UM1 AI144371 to B.F.H., P01 AI31251 to G.M.S., U01 AI169587 and R01 AI167653 to A.H, and NIH/NIAID Contract No. 75N93022D00003 to T.D. For data collection on a Titan Krios microscope, we utilized the cryo-EM facility at the Duke University Shared Materials Instrumentation Facility (SMIF), a member of the North Carolina Research Triangle Nanotechnology Network (RTNN), which is supported by the National Science Foundation (award number ECCS-2025064) as part of the National Nanotechnology Coordinated Infrastructure (NNCI). For processing data from the Titan Krios, this study utilized the computational resources offered by Duke Research Computing (http://rc.duke.edu; NIH 1S10OD018164-01) at Duke University. We thank Nilakshee Bhattacharya for assistance with Krios operations, Jordan Cocchiaro and Aaron Cook for program management support, Material and Structural Analysis Division of Thermo Fisher Scientific for operational support, and Duke School of Medicine for facilities and infrastructure support.

## RESOURCE AVAILABILITY

### Lead Contact

Further information and requests for resources and reagents should be directed to and will be fulfilled by the Lead Contact, Priyamvada Acharya.

### Materials Availability

Further information and requests for resources and reagents should be directed to Priyamvada Acharya. Plasmids generated in this study have been deposited to Addgene with the following accession numbers: 249831, 249832, 249833

### Data and Code Availability

- Cryo-EM reconstructions and atomic models generated during this study are available at wwPDB and EMBD (https://www.rcsb.org; http://emsearch.rutgers.edu) under the accession codes PDB IDs 9Y54, 9Y6H, 9Y6G, 9Y5L, 9YPD, 9YME and EMDB IDs 72507, 72562, 72561, 72512, 73294, 73104, 72966, 72943. The PDB ID 9MNH, 9MQN, 9NZ2 and EMDB IDs 48423, 48535, 49948 were already published previously. (Henipaviruses)
- This paper does not report original code.
- Any additional information required to reanalyze the data reported in this paper is available from the lead contact upon request.

## METHOD DETAILS

### Plasmids

All the proteins used in this study were recombinantly expressed by cloning relevant DNA sequences into pαH vectors with codon optimization for expression in human cells. All the constructs were synthesized by GeneImmune Inc. The OC43 spike proteins and were expressed with fusion of C-terminal T4 fibritin trimerization motif followed by HRV3C protease cleavage site, 8X Histidine tag and Twin-Step tag.

### Cell culture and transfection

The OC43 spike ectodomains, HNV protein and HIV envelopes were recombinantly expressed in Gibco FreeStyle 293F cells (Catalog no: R79007), whereas antibodies were expressed in Gibco Expi293F cells (Catalog no: A14527). The Gibco FreeStyle 293F and Expi293F cells, maintained at 37 °C and 8 % CO_2_ with shaking at 125 rpm were transfected at a viability of 95-100 % and cell density of 2 million cells per ml and 2.5 million cells per ml respectively. The separate transfections of cells with the plasmids encoding for S-OC43, HNV proteins (750 µg of plasmid per 1 liter of cells and HIV envelopes (600 µg of plasmid encoding HIV trimer and 150 µg of plasmid encoding furin) were mediated by Turbo293 transfection reagent (750 μg of plasmid per 1 liter of cells) from Speed Biosystems (Catalog No: PXX1002). For enhancement of the expression HyClone CDM4HEK293 media (Cytiva, Catalog No: SH30521.02) was added the next day after the transfection. The antibodies’ heavy and light chain plasmids were co-transfected into Expi293F cells using ThermoFisher Sci. ExpiFectamine kit (Catalog no A14524). ExpiFectamine kit enhancers were added approximately 16-18 hours after transfection, and cultures were incubated for 5 days. The medium containing the 293F and Expi293F cells was centrifuged at 4000 xg for 30 minutes at 22 °C to harvest the supernatant.

### Purification of coronavirus spikes proteins and HNV proteins

The supernatant was passed through a gravity column consisting of 10 mL of the Strep-Tactin resin (IBA LifeSciences, Catalog No: 2-1201) pre-equilibrated with 2 column volumes of Buffer W(1X) (IBA LifeSciences, Catalog No: 2-1003). After that column was washed with Buffer W(1X) until no absorbance was detected at 280 nm. The proteins bound to the column were eluted with the Buffer E(1X) (IBA LifeSciences, Catalog No: 2-1000) consisting of desthiobiotin and 25 mL fractions were collected. The fractions that contain proteins were identified in combination of absorbance at 280 nm using Thermo Scientific™ NanoDrop™ One^C^ Spectrophotometer and nano-DSF using the Tycho NT.6 from Nanotemper. The pooled fractions were concentrated and subjected to Size Exclusion Chromatography using Superose 6 Increase 10/300 (Cytiva, Catalog #: 29091596) at room temperature for further purification.

### Purification of HIV envelope

HIV envelope was purified using a Protein A column with immobilized PGT145 antibody. The PGT145 antibody was immobilized by cross linking using Disuccinimidyl Suberate (DSS). The column was maintained at pH 7.2 by rinsing with 1X PBS buffer. The supernatant that consists of the envelopes was passed through the column and washed with 5 to 6 column volumes of 1XPBS until the absorbance at 280 nm wavelength was zero. The envelope molecules bound to the column were eluted with 3M MgCl_2_. Post purification the column was rinsed with IgG elution buffer (100 mM Glycine, pH 3) and 1X PBS with 0.02 % NaN_3_ to have the pH at 7.2. The elution fractions were pooled and subjected to size exclusion chromatography using Superose 6 Increase 10/300 at room temperature for further purification.

### Purification of Antibodies

Purification of IgGs was carried out by using the Protein A columns (Thermo Fisher Scientific, Catalog #: 20366). The Protein A column loaded with supernatant was washed with to the column and washed with wash Buffer 1X PBS pH 7.5, until the absorbance at 280 nm reached baseline. The IgGs bound to the column were eluted with IgG elution buffer (100 mM Glycine, pH 3). The Fab fragments were generated by digestion with Endoproteinase Lys-C (Millipore SigmaCatalog no 11047825001). Both IgG and Fab were further purified by size exclusion chromatography to further purify using a Superose 6 Increase 10/300 column.

### Liposome Production

#### mRNA in vitro transcription and purification

Firefly luciferase mRNA was prepared by in vitro transcription (IVT) from a linearized plasmid template containing a T7 promoter, TEV 5′ UTR, a codon-optimized firefly luciferase CDS, and an *Xenopus laevis* β-globin 3′ UTR, with an encoded 101-A poly(A) tail (Pardi et al. 2015). Briefly, Plasmid DNA was linearized and IVT was performed at 37 °C with Promega T7 Enzyme Express Mix, a standard rNTP mix (ATP, CTP, GTP), and N1-methylpseudouridine triphosphate (TriLink Biotechnologies, N-1081) to fully substitute uridine residues. Co-transcriptional capping was performed with CleanCap AG 3′OMe (TriLink Biotechnologies, N-7413). Following transcription, residual plasmid DNA was digested with DNase I (NEB, G8002B) and quenched with 0.5 M EDTA (Promega, V4231). Crude IVT material was purified using a multi-step downstream workflow consisting of tangential flow filtration (TFF) with a 100 kDa mPES cassette (Repligen, PP100LP2L) for buffer exchange into HEPES/NaCl/EDTA. Multimodal chromatography was used to purify the mRNA to remove rNTPs and DNA fragments, followed by a second TFF polishing step into citrate buffer pH 6.4. RNA was sterile-filtered with a 0.2 µm PES filter (Supor membrane, Cytiva), and stored at −80 °C (Whitley et al. 2022).

#### Liposome Preparation and Purification

Cationic liposomes were prepared using an in-house formulation (Avanti Polar Lipids: POEPC, 1,2-dioleoyl-sn-glycero-3-ethylphosphocholine, Cat. 890704; DOPC, 1,2-dioleoyl-sn-glycero-3-phosphocholine, Cat. 850375; cholesterol, Cat. 700000; and DSPE-PEG2000, MedChemExpress, Cat. HY-140739) at 65:10:20:5 molar ratio. Purified in vitro transcribed mRNA was diluted to 0.2 mg/mL in 50 mM citrate buffer, pH 4.0, immediately prior to encapsulation. Liposome synthesis was performed using the Knauer IJM-based Nanoscaler system (Knauer Wissenschaftliche Geräte GmbH) at a mixing ratio of 3:1 aqueous to solvent volumetric ratio following manufacturer specifications. Empty liposomes were produced by loading the aqueous phase loop with 50mM citrate buffer, pH 4.0, instead of mRNA. The liposomes were collected from the Nanoscaler outlet and immediately concentrated and buffer exchanged using Amicon Ultra centrifugal filters (100 kDa cutoff, Millipore Sigma, Cat. UFC910024). Briefly, filters were sanitized with 0.5 N sodium hydroxide for 10 minutes and rinsed twice with PBS. Samples were concentrated to approximately 500 μL by centrifugation at 4 °C for 10 minutes at 1000 x*g*, diluted 5-fold in PBS, and concentrated again by centrifugation at 4 °C for 10 minutes at 1000 x*g*. RNA encapsulation efficiency was quantified using the Quant-iT RiboGreen assay (Thermo Fisher Scientific, Cat. R11490) following manufacturer protocol on a SpectraMax iD3 microplate reader (Molecular Devices). Total RNA content was measured after addition of 0.1% Triton X-100 (Sigma Aldrich, Cat. X100) to fully disrupt liposomes. Encapsulation efficiency was calculated as a percentage of encapsulated RNA divided by total RNA. Final liposome formulations were diluted with 30% Sucrose for cryoprotection and stored at −80 °C until processing.

### Expression and production of an HIV-1 Env-based ferritin nanoparticle

DNA constructs containing the HIV-1 Env-based ferritin nanoparticles were synthesized (Genscript), produced and purified as previously described (Saunders et al. 2024). Plasmids encoding the ferritin nanoparticles were transiently transfected in FreeStyle 293 cells (ThermoFisher) using 293Fectin (ThermoFisher). All cells were tested monthly for mycoplasma. The construct contained an HRV3C-cleavable C-terminal twinStrepTagII-8×His tag. On day 5, cell-free culture supernatant was generated by centrifugation of the culture and filtering through a 0.8-μm filter (Nalgene). Protein was affinity purified from filtered cell culture supernatants using a StrepTactinXT column connected to an AKTApure FPLC system (Cytiva). The nanoparticle was incubated with 10 % (wt/wt) HRV3C protease for 16 hours at room temperature. Cleaved protein was then passed over NiNTA resin to remove cleaved tags before being run over a HiLoad 16/600 Superose6 column (Cytiva) in 10 mM Tris pH 8.0, 500 mM NaCl. The protein fractions were pooled, concentrated using a Centricon70 (Millipore), filtered using a 0.22 μm filter (Millipore), and snap frozen for long-term storage at −80 °C.

### Production of EABR eVLPs

EABR eVLP was produced following methods as described (Hoffmann et al. 2023; Shoemaker et al. 2025). HIV-1 Env EABR eVLP was expressed by co-transfecting Expi293F cells with a furin expressing plasmid at 4:1 ratio. Expressed cells were allowed to grow for 65-70 hours on an orbital shaker at 37 °C and 8% CO_2_ before harvesting by spinning down the cells at 4000 xg for 15 minutes. Supernatant containing EABR eVLP was collected and filtered with 0.2 µm filter unit before sucrose cushion centrifugation. A 4 ml volume of 20% w/v sucrose PBS was added to centrifuge bottles and 20 ml of filtered supernatant was layered on top of it followed by ultracentrifugation using Optima MAX-XP (Beckman Coulter) at 4 °C Optima MAX-XP for 2 hours at 48,000 rpm (rcf average = 150,700). After ultracentrifugation, the supernatant was removed and 250 μL of sterile 1X PBS + 0.02% NaN_3_ was added to EABR eVLP containing pellet without resuspending. The pellet was then allowed to be incubated in 4°C overnight before pipetting up and down to resuspend. The resuspended pellet was then incubated at 37 °C for 15 minutes followed by spinning down at 11,000 xg for 2 minutes. Supernatant was collected and was further spun down for 3 times to remove residual cell debris to be further purified by SEC using a Superose 6 Increase 10/300 column (Cytiva) equilibrated with 1X PBS + 0.02 % NaN_3_. Fractions eluted in the void volume were collected and combined for Cryo-EM grids freezing.

### H5N1 Influenza virus

Allantoic fluid collected from H5N1 (A/Texas/37/24) infected eggs was inactivated in 4 % methanol-free formaldehyde (Thermo scientific Cat. 047340.9M) for at least 30 minutes at BSL3 containment. Inactivated virus was then processed at BSL2 for downstream analysis. In short, inactivated virus was layered with a 30 % sucrose cushion and then concentrated via ultracentrifugation (Sorvall wX+ Ultra Series centrifuge) at 27,500 rpm for 1 hour at 4 °C. After centrifugation, sample was resuspended in 100ul PBS and stored at 4 °C until use.

Prior to imaging, the sample was mixed with 0.085 mM DDM.

### Grid Freezing

The spike samples were mixed with 0.5 % Glycerol, LayV samples with 2 % Glycerol and AngV samples with 0.005 % n-dodecyl β-D-maltoside (DDM) before plunge freezing. HIV-1 Env samples were mixed with 0.005 % DDM before plunge freezing.

The S-OC43 samples were incubated with excess amount of Fabs in 1:6 ratio and S-HKU1 samples with 1:4:4 ratio for 30 minutes and mixed with 0.5 % glycerol right before freezing the sample. The HIV envelope with was incubated with the vaccine-elicited antibody at 1:5 ratio, followed by mixing with 0.005 % DDM prior to plunge freezing. The Langya virus fusion protein was incubated with the 22F5 Fab at 1: 4.87 ratio and mixed with 2 % glycerol for freezing.

For all the samples, the Quantifoil R1.2/1.3 Cu300 (Electron Microscopy Sciences, Catalog no: Q350CR-14) grids were glow discharged at 15 mA for 15 seconds using PELCO easiGlow™ from TED PELLA, INC. The EM GP2 Automatic Plunge Freezer from Leica Microsystems was used for applying sample, blotting, plunging and vitrification. Sample blotting was carried out at 22 °C under 95 % humidity. A volume of 2.5 - 3µL sample with 1- 4.5 mg/mL concentration was applied on to the grids and blotted with a 30 seconds delay followed by 2.5 seconds blotting. The samples were plunged into liquid ethane and vitrified in liquid nitrogen.

### Data Collection and Processing

The data were collected on 100 kV TFS Tundra and 300 kV TFS Krios microscopes with different detectors as mentioned in Table1. The collected micrographs were subjected to Patch motion correction, Patch CTF estimation for phase flipping and amplitude correction using the CryoSPARC software. Blob picker, extraction from micrographs, 2D classification, Select 2D classes were followed as mentioned in the Figure 3 B. The 3D reconstruction of the maps was carried out using ab-initio reconstruction in CryoSPARC followed by multiple refinements including the hetero refinements, Non-uniform refinements and the local refinements. A structure of S-OC43 spike available in Protein Data Bank (PDB), PDB ID:6OWH, was used as an initial model for fitting into the density maps. The refinements were carried out utilizing Coot, Phenix, Chimera and Pymol in combination. (Pettersen et al. 2021; Emsley et al. 2010; Adams et al. 2010).

